# Decoding the oxytocinergic and behavioral signatures of milk ejection

**DOI:** 10.64898/2026.07.12.738011

**Authors:** Wei Xiao, Qian Zheng, Yang Wang, Ying Yuan, Yuge Chen, Tianxing Zheng, Yuzhu Chen, Yijia Gao, Bingrui Song, Bin Zhang, Liyao Qiu, Linghui Zeng, Huan Ma, Colin H Brown, Shumin Duan, Gang Pan, Zhihua Gao

**Author notes:** Lead Contact: Zhihua Gao. These authors contributed equally to this work (Wei Xiao, Qian Zheng).

## Abstract

Oxytocin-mediated milk ejection (ME) is pivotal to effective breastfeeding and productive health, yet behaviorally decoding and revealing neural mechanisms of ME remains challenging. Here, we combined *in vivo* calcium imaging and intramammary pressure recording to uncover the temporal connections between episodic activity of oxytocin neurons and ME in conscious lactating rats. Leveraging the association and behavioral responses in dam and pup, we developed a supervised machine learning framework (ME Decoder) to enable automated analyses of ME. Inspired by its interpretable features, we defined the activity-coupled dam-pup interactions (ADPI), manifested by high kyphosis of the dam followed by pup treading and stretch, as the behavioral signatures of ME. By ME Decoder and ADPI analyses, we detected reduced ME but unaffected activity of oxytocinergic neurons after systemic blockade of oxytocin receptor. Our study uncovers the oxytocinergic and behavioral signatures of ME and provides a generalizable approach for further investigation.

## Introduction

Breastfeeding is a universal behavioral feature of mammals that ensures the survival of the infants, fosters mother-infant bonding and supports the propagation of the species [1]. Central to lactation is the suckling-induced milk ejection, a neuroendocrine reflex driven by the pulsatile release of oxytocin from hypothalamic neurons into the bloodstream [2–4]. Circulating oxytocin then acts on the myoepithelial cells of the mammary gland to induce contraction of the alveoli and drive milk ejection [5, 6].

Early studies revealed that hypothalamic oxytocin neurons intermittently increase their action potential firing rate by 20-40-fold during suckling [7, 8]. These bursts occur at the same time across the oxytocin neuron population [9, 10], which generates pulsatile oxytocin release to triggers milk ejection [11, 12]. Using genetic labeling and *in vivo* calcium imaging, recent studies have reported pulsatile activity of oxytocin neurons in the paraventricular nuclei (PVN) during lactation in conscious animals [13–17], reminiscent of the milk ejection bursts observed previously [7, 8]. However, lactation is a dynamic and reciprocal process between the mother and offspring that extends beyond nutrition to shape maternal-infant bonding and maternal psychological wellbeing [18, 19]. Predominant studies focus on changes in maternal oxytocin neuron activity but miss the essential behavioral features of milk ejections [13, 14, 17]. A comprehensive understanding of lactation requires the integration of both maternal oxytocinergic signals and the behaviors of the dam and pups.

Computational ethology quantifies animal behaviors by computer vision and machine learning, and provides an objective framework different from traditional manual scoring [20, 21]. Deep learning has enabled precise tracking and classification of complex behaviors across species [22–24]. Skeleton-based approaches(e.g. DeepLabCut or SLEAP [25–26]) offer greater interpretability but are limited in three-dimensional motion, occlusions, and multi-animal interactions [24]. By contrast, image-based approaches (e.g. DeepEthogram [27]) capture rich spatiotemporal information, yet sensitive to background variability and computational resources [28]. Breastfeeding involves close physical interactions between the dam and pups, during which frequent occlusions and complex motions occur. Thus, reliably detecting milk ejections demands a framework with computational robustness and physiological interpretability.

To address these issues, we combined *in vivo* recoding of oxytocinergic activity with intramammary pressure (IMP) measurements and ethological analyses to characterize the neural and behavioral features of milk ejection in lactating rats. Our study defines the oxytocinergic and behavioral signatures of milk ejection and provides a foundation for future investigation.

## Methods

### Animals

Sprague Dawley (SD) and OXT-Cre rats were used in the experiments. Generation of OXT-Cre rats has been previously described [29]. Animals were maintained and group-housed under 12 h light/dark cycle (lights on at 7:00 am) at 24±2°C (humidity 40 %-60 %), with water and food ad libitum in the Laboratory Animal Center of Zhejiang University. All animal procedures were conducted in accordance with protocols approved by the Committee of the Ethics of Animal Experiments of Zhejiang University.

### Stereotaxic injection

Female OXT-Cre/SD rats were anesthetized with 45 mg/kg sodium pentobarbital and placed on a heating pad to maintain body temperature. Rats were head-fixed in a stereotaxic instrument (RWD, Shenzhen, China) and viruses were injected using a 2.5 μL Hamilton syringe driven by a micro-injection pump (KD Scientific). Viruses were slowly infused into the targeted brain regions for at least 10 min and the syringe was maintained *in situ* for another 10 min after viral infusion. To avoid viral backflow, the syringe was held for 1 min after withdrawing 50 μm before the complete withdrawal.

For calcium recording of oxytocin neurons, the AAV2/9-DIO-hSyn-GCaMP6m-WPRE-pA (1.00 x 10^13^ genomic copies/mL, 150-300 nL) were injected into the SON (AP, -1.35 mm; ML, +2.10 mm; DV, -9.5 mm) in female OXT-Cre, 5 days before mating. A 400 μm optical fiber (Inper, Hangzhou, China) was implanted 200-300 μm above the virus injection site. For central drug administration, a cannula (OD, 0.48 mm; ID, 0.34mm; 622003, RWD, Shenzhen, China) was implanted into the lateral ventricle (AP, -0.96 mm; ML, +2.0mm; DV, -4.2mm) in female SD rats during the same surgical session as virus/fiber implantation.

### Immunofluorescence analysis

Rats were anesthetized with 60 mg/kg sodium pentobarbital and perfused with normal saline followed by 4% paraformaldehyde (PFA, Sigma-Aldrich). Brains were dissected, post-fixed in 4% PFA overnight, and transferred to 30% sucrose in 0.1 M phosphate buffer solution (PBS) for 2 days. Brains were embedded in OCT ( SAKURA Tissue-Tek) and coronal sections were cut at a thickness of 30 μm using a microtome (NX50, Thermo Fisher Scientific). Antigen retrieval was carried out as described previously [29]. To label oxytocin neurons, slices were incubated with OXT-neurophysin antibody (Mouse, 1:3000, Millipore) in 5% BSA overnight at 4°C, washed in Tris-buffered saline (TBS), followed by secondary antibody incubation for 2 hours at room temperature. Nuclei were counterstained with 6-diamidio-2-phenylindole (DAPI) and slides were mounted with antifade reagents (P10144, Invitrogen). The immunofluorescence images were captured by FV-1200 confocal microscope (Olympus) or VS120 microscope (Olympus) and cell number was counted using the ImageJ software (NIH).

### Preparation of lactating animals and video recording

Female OXT-Cre/SD rats aged 12-14 weeks (240 g-280 g) were mated with stud male SD rats. Rats at mid-pregnancy were gently handled daily until parturition. Immediately after parturition, the number of pups was adjusted to 8, including 4 males and 4 females per litter, as previously described [7, 12]. The dams and pups were habituated in a black acrylic test arena for 30 min, one day before the experiments. For lactation-related recording [30, 31], dams were separated from their pups for 4 h, then introduced to the test arena and reunited with pups for lactation. For systemic oxytocin receptor antagonism experiments, dams were separated from their pups for 30 min before injection of 5 mg/kg L-368,899 (HY-108677, MedChemExpress) or 0.9% saline and the pups returned 30 min later for recording. For the activation and inhibition of central oxytocin receptors, dams were separated from their pups for 30 min before injection of L-368,899 (25 mM, HY-108677, MedChemExpress) or artificial cerebrospinal fluid injected, and the pups returned 15 min later for recording. The process was videotaped by an overhead RGB camera with a resolution of 640×480 pixels for 0.5-1 h, in a 45 × 45 × 40 cm (width × length × height) arena with fresh bedding and 40 Lux illumination.

### Fiber photometry recording and data analyses

The GCamp6m fluorescence signals were recorded using a dual-color recording system (Inper, Hangzhou, China), equipped with a 470 nm channel to excite GCaMP6 and a 410 nm channel as control. The activity of oxytocin neurons was recorded in dams during lactation (either awake or anesthetized) from postpartum day 5-18 for interactions with their pups, including pup retrieval and crouching. The power output at the tip of the fiber was ∼30 μW and signals were recorded at a frequency of 50 Hz.

Data were analyzed using customized software provided by the manufacturer and homemade MATLAB code. Processed signals were derived by smoothing the raw data with a 10-point sliding window average, followed by detrending. Candidate calcium transients were automatically detected from these signals using the MATLAB *findpeaks* function, applying a minimum prominence threshold of 50% of the maximum signal and a minimum FWHM of 1.5 s [13, 32]. To distinguish true oxytocin neuron calcium transients from noise and motion artifacts, a peak was considered valid only if it fulfilled the following criteria: its amplitude exceeded twice the standard deviation (2×SD) of the baseline (calculated from -10 to -5 s pre-peak), and no correlating deflection was observed in the 410 nm channel. The onset of each validated peak was then defined as the time point at which the signal first rose to 10% of the peak amplitude, using the mean of the -10 to -5 s pre-peak interval as baseline [33]

For the kinetic analyses of calcium signals, we extracted the smoothed data from -5 s to +15 s around the rise of peak. The ΔF/F was calculated as (F-F0)/F0, with F0 representing the average fluorescence of the baseline (-5 s to 0 s). To compare the calcium signals across different animals and situations, the ΔF/F data were converted into Z-scores [34, 35]. The area under curve (AUC) was calculated by trapezoidal integration of ΔF/F signals relative to baseline mean during the 0-5 s response window, and the peak half-width was defined as the time interval between 50% rise and 50% fall points.

### Intramammary pressure measurement

IMP was measured as previously described [36], along with simultaneous calcium recording from oxytocin neurons. Prior to the procedure, dams were separated from their pups for 18 h to engorge the mammary gland. Dams were anesthetized via i.p. injection of sodium pentobarbital (35 mg/kg) and placed supine on the operating area. The fur around the lower abdominal nipples was shaved and the area was disinfected with iodophor. Following blunt dissection of the mammary duct, a polyethylene catheter (outer diameter: 0.96 mm, inner diameter: 0.6 mm; RWD, Shenzhen, China) was inserted into the duct through a small incision. The catheter, pre-filled with 0.15 M sodium acetate, was connected to a portable blood pressure sensor (ADInstruments, Shanghai, China). IMP signals were amplified via a bridge amplifier (ADInstruments) and recorded in real-time using LabChart Pro software (ADInstruments). Post-cannulation, 6 pups were reintroduced for suckling, positioned to avoid direct contact with the cannulated nipples.

### Manual behavioral analysis of lactation videos

#### Behavioral analysis criteria

Lactation videos were manually annotated frame-by-frame using Elan (version 6.6) by two independent experimenters, who were both blind to animal identity and neural responses. There was high consistency (>90%) between annotations performed by different individuals.

During natural lactation, pups are largely concealed beneath the dam’s abdomen, making it challenging to observe behavioral changes across the entire litter, even when concurrently recorded from the top, side, and bottom. The top view was selected for subsequent analyses because it provided the best view of both dams and pups, maximizing the number of visible pups. Based on the behavioral sequences of unobstructed pups during suckling, we categorized their actions into four different types (representative examples shown in Fig. 5B): P0, quiescent suckling, pups latched to nipples in a relaxed posture without body movements; P1, treading, suckling pups pushing the nipples with their forelegs and treading with their hindlegs; P2, stretching, suckling pups in a rigid and fully extended posture; P3, switching, pups temporarily detach from one nipple and relocate to another.

For lactating dams, we further classified their nursing patterns into four types: (representative example illustrated in Fig. 5B): D0, low kyphosis crouching, crouching dam characterized by a hunched posture with minimal spinal curvature crouching over pups; D1, High kyphosis, crouching dam marked by a pronounced arched back posture; D2, cleaning, crouching dam with self-grooming or pup-grooming; D3, adjusting, dam with brief postural shifts including repositioning or transiently quitting crouching.

#### Annotated data analysis

Annotated videos in .eaf files were automatically converted into binary behavioral matrices by Python code (detailed in GitHub) with a temporal resolution of 1 s, where 1 denotes the occurrence of a specific behavior and 0 denotes its absence. These matrices were then used for subsequent behavioral quantification and sequence analysis. To investigate temporal patterns during milk ejections, we extracted behavioral matrices for dams and pups within a 25-second time window following the onset of calcium waves (77 trials total). For non-milk ejection events, a 25-second window was randomly selected from suckling periods excluding milk ejections (231 trials total). We computed the probability of each behavior’s occurrence relative to oxytocin neuron calcium waves. The resulting data were organized into a 25×4 matrix, with columns representing behavioral categories and rows denoting time-dependent probabilities. Finally, these matrices were uploaded to the OEBiotech Cloud Platform (https://cloud.oebiotech.com) to generate Circos plots visualizing temporal behavioral dynamics.

#### Criterion for activity-coupled dam-pup interactions

To establish a reliable reference to recognize milk ejections in the learning process, we defined ADPI into four phases: the silent, active, consummatory, and energized phases, according to the temporal responses of dam and pups. Typically, the dam crouched over pups in a low kyphosis with pups quietly suckling. This phase, characterized by minimal activity, was referred to as the silent phase. Later, the dam moved into high kyphosis with pups treading legs, this was considered as the start of ADPI, the active phase. In the consummatory phase, the pups engulfed a large amount of milk and exhibited an extended stretch posture, indicating the climax of ADPI. After the extension, pups showed variable activities, with some directly changing into the silent phase, some still treading actively, some rotating their bodies, and some detaching from nipples. This was considered as the end of ADPI, the energized phase. After the above four phases, the dam and all pups would revert back into the silent phase.

#### Design of ME Decoder

To detect milk ejections (ME) in videos, we developed ME Decoder, a two-stage framework inspired by temporal action localization methods [37]. ME Decoder has two modules: the ME Detector and the ME Refiner. The ME Detector module comprises a Temporal Shift Module (TSM) and a Boundary Matching Network (BMN) [38, 39]. The ME Refiner module also consists of the TSM for classification.

In detail, the TSM in ME Detector takes a query video as input and outputs behavioral representation for each frame. Then, the BMN uses TSM-generated behavioral representations to locate candidate ME segments. These segments are fed into the TSM in the ME Refiner for classification as genuine ME segments, or not.

#### Data analysis related to ME Decoder

All data analyzed related to ME Decoder, including the IoA, IoU, milk ejection durations, intervals correlated with calcium activity, model contributions and behavioral action contributions, were documented and implemented in the code.

#### Quantification and statistical analysis

All measurements were exported to GraphPad Prism 9 and analyzed by Student’s t test, one-way ANOVA or mixed effect model depending on the structure of the data. Nonparametric tests were used if the data did not match assumed Gaussian distribution. Data are presented as Mean ± SEM, with statistical significance taken as \**p* < 0.05, \*\**p* < 0.01 and \*\*\**p* < 0.001. Briefly, for behavioral analysis and model evaluation, Student’s t test was used, with n = 5 - 9 rats or n = 40 - 70 trials, as indicated in the figure legends. For calcium signal comparison, one-way ANOVA or mixed effect model were used as appropriate.

## Results

### Maternal oxytocin neurons exhibit episodic calcium activity during continuous suckling

To uncover the activity of magnocellular oxytocin neurons during lactation, we recorded the calcium responses of oxytocin neurons in the SON using the calcium sensor, GCaMP6m, in conscious rats in mid-lactation (postpartum day 11, PPD11) (Fig 1A-B and Fig S1A). While oxytocin has been shown to regulate maternal behavior [40–43], we observed no changes in calcium signal in oxytocin neurons when dams retrieved or groomed pups (Fig 1C and Fig S1B-C), likely due to the low sensitivity of calcium recording. Neither did the calcium signal change at the dam-pup reunion or at the onset of suckling (Fig 1C–1D and Fig S1D). After continuous suckling for several minutes (661.2 ± 78.6 s), maternal oxytocin neurons exhibited a dramatic calcium elevation (Fig 1C and 1E), which was much larger than the small transients previously observed during social touch [44]. Such robust calcium elevations occurred every few minutes subsequently during suckling (Fig 1F), with a mean interval of 318.2 s (318.2 ± 48.1 s), a duration of 5.9 s (5.9 ± 0.2 s), a rapid rise to peak within 2 s, and a return to basal levels within 4 s (Fig 1E and Fig S1E-H). These episodic calcium waves are temporally consistent with the time course and duration of burst firing of oxytocin neurons during electrophysiological recordings in lactating rats [12], as well as with the calcium waves in PVN oxytocin neurons of lactating mice and rats [13–17].

**Fig 1.**
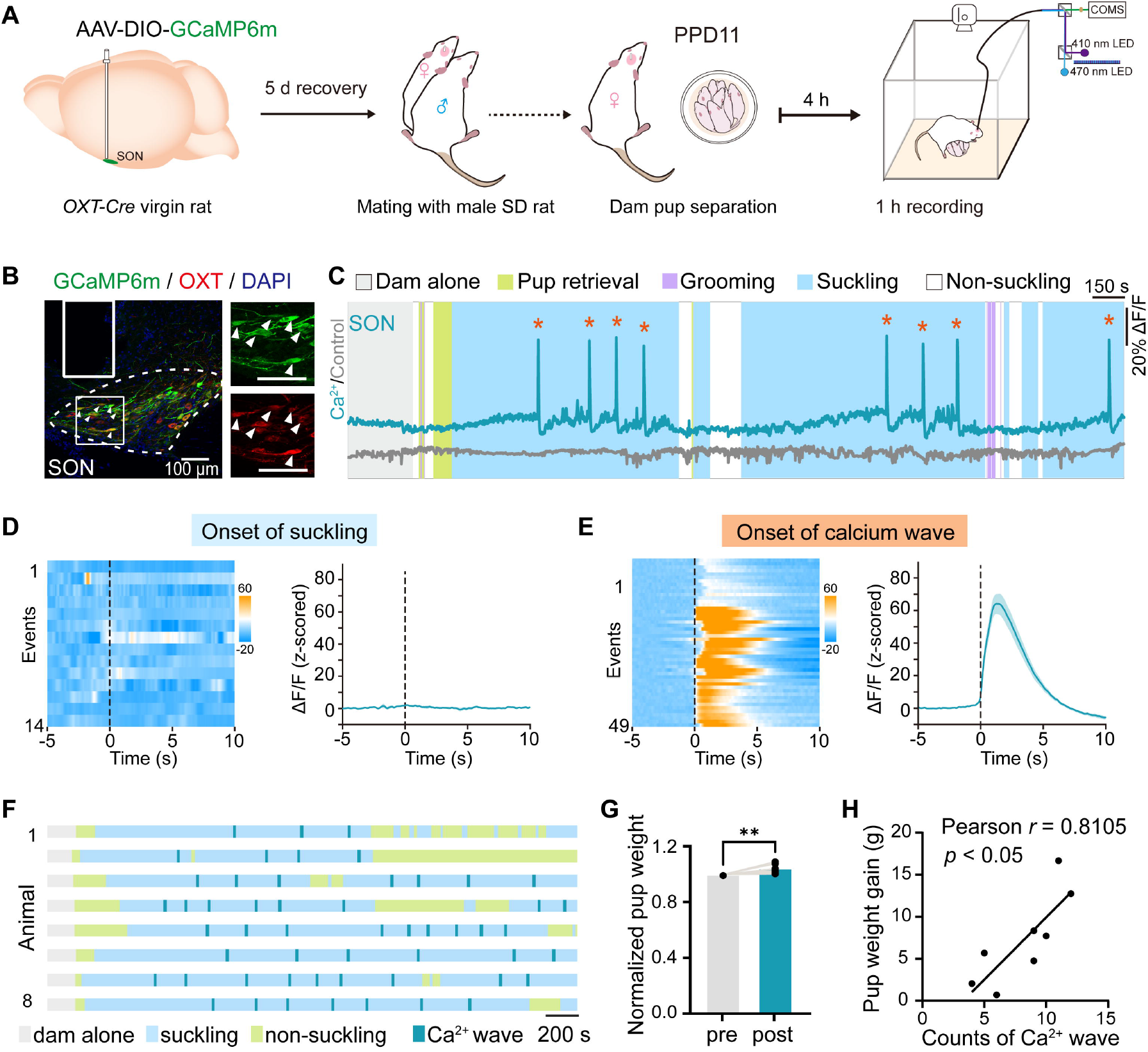
Maternal oxytocin neurons exhibit episodic calcium activity during continuous suckling. **(A)** Schematics of experimental procedure and fiber photometry recording setup. **(B)** Virus expression and oxytocin staining showing the co-labeling of GCaMP6m (green) and oxytocin neurons (red) in SON. **(C)** Representative traces of the calcium (470 nm, blue) and control signals (410 nm, gray) showing the pulsatile activities of oxytocin neurons in SON during suckling. Asterisks indicate calcium waves. **(D)** Individual (left, heatmap) and average (right, mean ± SEM) calcium signals aligned to the onset suckling (n = 5 dams, 14 trials). **(E)** Individual (left, heatmap) and average (right, mean ± SEM) calcium waves during suckling (n = 5 dams, 49 trials). **(F)** Raster plots displaying the occurrence of calcium waves during suckling in 1 h recording (n = 8 dams). **(G)** Pup weight (8 pups per litter) measured before and after 1 h lactation (n = 8 litters). ** *p* < 0.01, two-tailed paired t-test **(H)** Pearson correlation analyses between calcium waves and pup weight gain (n = 8 litters). r = 0.8105, *p* < 0.05

Similar to previous observations in mice [13, 17], episodic calcium waves discontinued when suckling stopped (Fig 1F). Moreover, local application of lidocaine to inhibit sensory transmission from nipples [45], significantly delayed the onset of calcium waves and reduced the number of calcium waves measured over 1 h (Fig S1I-K), suggesting that pup suckling triggers the pulsatile activity of oxytocin neurons through sensory inputs around the nipples. In addition, the number of calcium waves positively correlated with the weight gain of pups over 1 h (Fig 1G–1H), indicating tight associations between calcium elevations in oxytocin neurons and milk ejection.

### Attenuated suckling-evoked oxytocin neuron activity and milk ejection during pentobarbital anesthesia

To examine the temporal correlation between oxytocin neuron activity and milk ejection, we simultaneously recorded calcium and IMP in anesthetized lactating dams. Under pentobarbital anesthesia, the first calcium wave of oxytocin neurons occurred ∼40 min after suckling onset, in contrast to ∼10 min in conscious dams (Fig 2A–2B). Overall, the calcium waves in anesthetized dams had a significantly smaller amplitude, shortened duration, decreased area under curve, and slightly longer inter-wave interval (Fig 2C-I). Using concurrent IMP recording, we observed that each calcium wave was followed by an IMP rise (Fig 2J), with a delay of 15.0 ± 2.9 s (Fig 2K), consistent with the latency between the burst firing of oxytocin neurons and IMP elevation reported previously [7, 12]. These data suggest that episodic calcium waves represent the synchronized bursts of oxytocin neuron population, which induce milk ejection.

**Fig 2.**
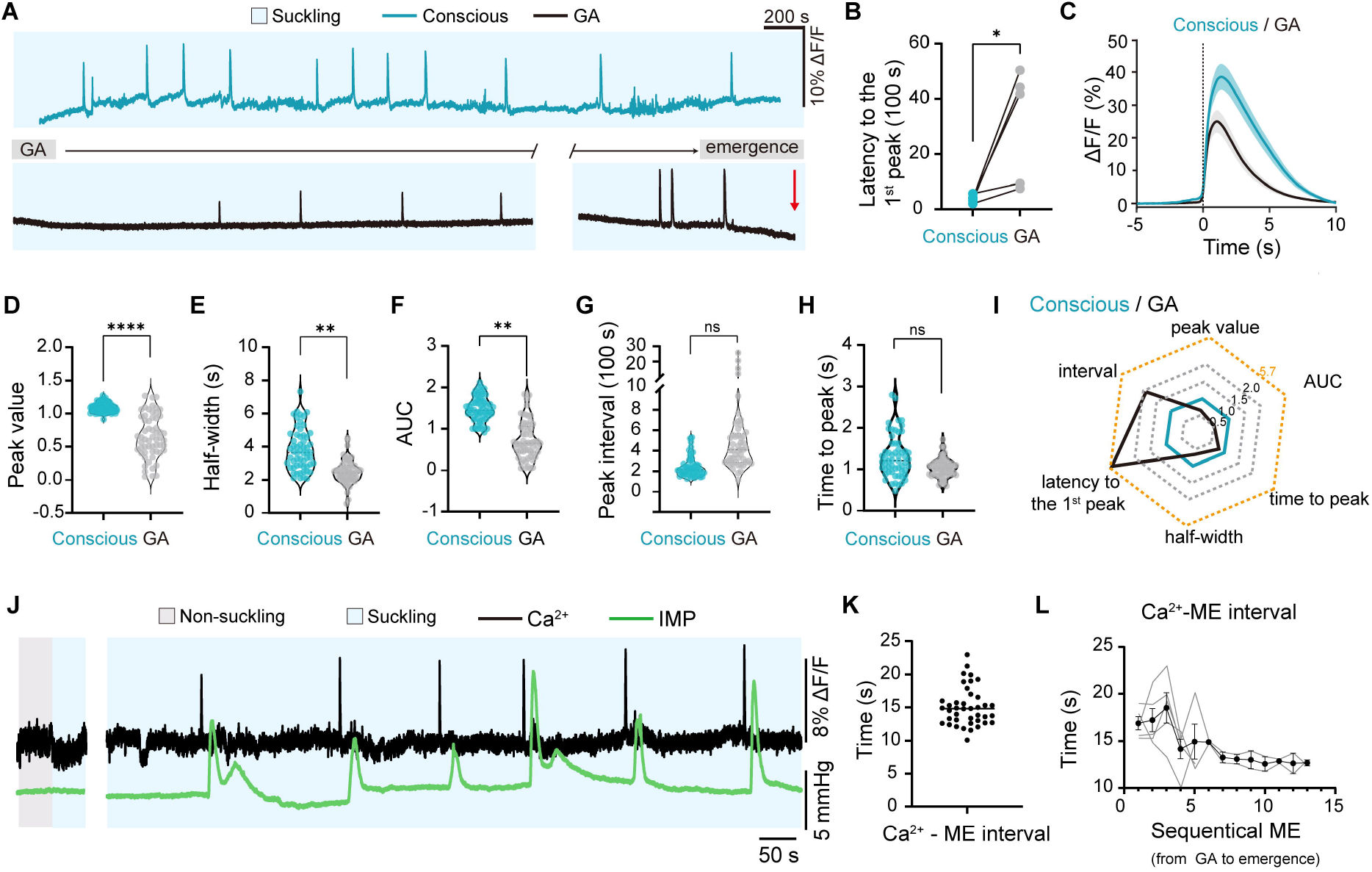
Anesthesia inhibits the activity of oxytocin neuron population and milk ejection. **(A)** Representative traces showing calcium signals of oxytocin neurons during suckling in conscious and anesthetized dams.**(B)** Quantification of the latency to the first calcium peak during suckling (n = 5 dams). * *p* < 0.05, two-tailed paired t-test **(C)** Average (mean ± SEM) calcium waves of oxytocin neurons during suckling, aligned to the onset of calcium wave, in conscious and anaesthetized dams (n = 5 dams, 58 trials). **(D-G)** Peak amplitude (D), half-width (E), area under curve (F), inter-peak intervals (G) and time to peak (H) of calcium waves during suckling in conscious and anesthetized dams. (n = 5 dams, 59 trials). *\*\* p* < 0.01, *\*\*\*\* p* < 0.0001, *p* = 0.9246 for G, *p* = 0.153 for H Mixed-effects model. **(I)** Radar chart visualizing multi-dimensional characteristics of calcium waves in oxytocin neurons during suckling in conscious versus anesthetized dams. **(J)** Representative traces of simultaneously recorded calcium signals of oxytocin neurons (black) and intramammary pressure (green) obtained from an anesthetized dam during suckling. **(K)** Quantification of the latency from calcium elevation to IMP rise during suckling (n = 5 dams). **(L)** Latency from calcium elevation to IMP rise during suckling, plotted in the sequential of occurrence of milk ejections (n = 5 dams). Gray lines represent individual animals; the dark line represents the mean ± SEM.

Notably, as animals gradually recovered from anesthesia, the calcium wave-to-IMP interval gradually shortened and stabilized at 13.6 ± 1.0 s by the time of emergence (Fig 2L), in line with the time course required for oxytocin to reach and affect the mammary gland following release form the posterior pituitary [12]. Together, these data demonstrate that pentobarbital anesthesia blunts suckling-induced activation of oxytocin neurons and peripheral oxytocin actions, preventing the full expression of the milk-ejection reflex.

### A deep learning framework automates milk ejection detection

The requirement for general anesthesia during IMP measurement precludes its use in assessing milk ejection in conscious animals. To investigate lactation-related behaviors and their health benefits [46, 47], it is important to develop an approach to enable noninvasively quantification of milk ejections in freely behaving dams. Previous studies have shown that pups may exhibit body extension (pup stretching) upon ingesting milk [31, 48, 49], suggesting that milk ejections can be decoded behaviorally. To test this possibility, we adopted a computational ethology approach, aiming to identify behavioral signatures of milk ejection under natural conditions.

We first applied DeepLabCut to track keypoints on both dams and pups. While the tool effectively tracked the dams, reliable pup tracking was unsuccessful due to their frequent occlusion by the dam. Keypoint analyses of the dam revealed transient displacements and angular changes in the skeleton of the lactating dam, but none of them correlate with the episodic occurrence of calcium waves (Fig 3A–3B). These data indicate that skeleton-based postural changes of the dam alone are insufficient to represent milk ejections.

**Fig 3.**
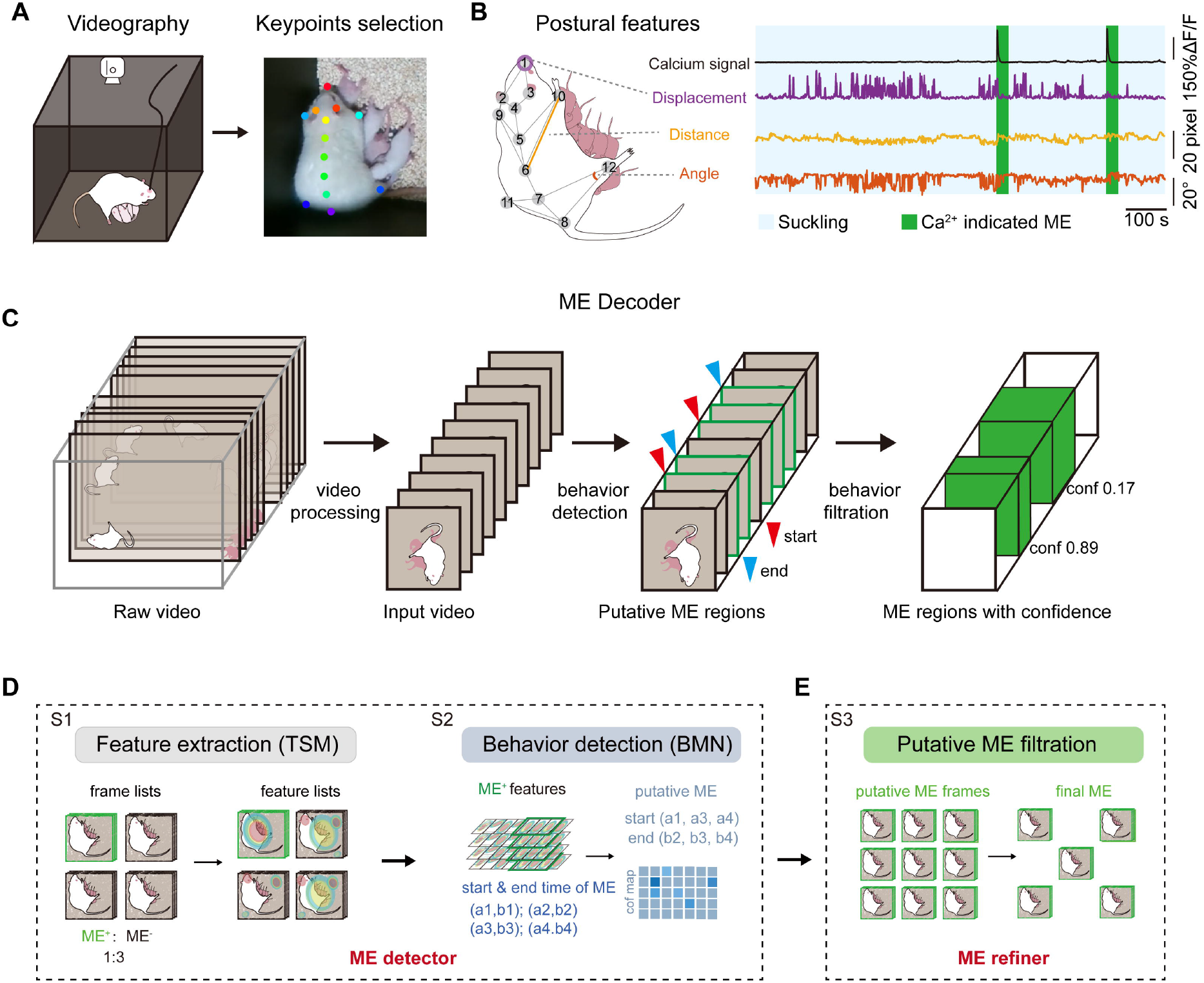
A deep learning framework automates milk ejection detection. **(A)** Schematic illustration of camera recording (left) and video frame with keypoints tracking (right). **(B)** Illustrations of postural features (left) and representative temporal dynamics of features across suckling and calcium waves (right). **(C)** Schematic illustration of the main structure in ME Decoder. **(D-E)** Schematic illustration of key steps of ME Decoder, including the ME detector (D) and ME refiner (E).

To circumvent the limitations of the skeleton-based approaches, we designed an image-based supervised framework, ME Decoder, to fully capture behavioral changes related to milk ejections (Fig 3C). ME Decoder integrates a Temporal Shift Module (TSM) to extract dynamic spatiotemporal features of the video [38], and a Boundary-Matching Network (BMN) to localize the onset and offset of the putative milk ejection episodes (Fig 3D) [50]. BMN-proposed candidate episodes are further filtered by a TSM-based classifier to minimize false positives (Fig 3E).

### Dam-pup interactions contribute to the recognition by ME Decoder

ME decoder was trained to recognize milk ejections according to the coupling time between episodic oxytocinergic calcium waves and milk ejections (Fig 2J-K). Video segments of 9 s duration, starting ∼10 s after the onset of calcium waves, were used as positives for model training, whereas 9 s video segments from the rest videos as negatives (Fig 4A). Following architectural optimization, ME Decoder reliably identified milk ejection episodes, achieving an F1 score over 0.8 with IoA (intersection over area) around 0.6 in the validation dataset (Fig 4B). When applied to a new test dataset (dataset 1, comprising 87 milk ejections in 9.58 h of videos), the model identified more than 75% of milk ejections, with ∼90% accuracy (Fig 4C). The majority of automatically identified milk ejection (Ai-ME) video segments showed durations of 8-10 s (Fig 4D), with 79.4% occurring 6-10 s after calcium waves of oxytocin neurons (Fig 4E). These data are highly overlapped with the ground truth milk ejections (GT-ME, defined as events occurring 7-16 s after calcium elevation) (Fig 4F-4G), suggesting that ME Decoder reliably detected milk ejections.

**Fig 4.**
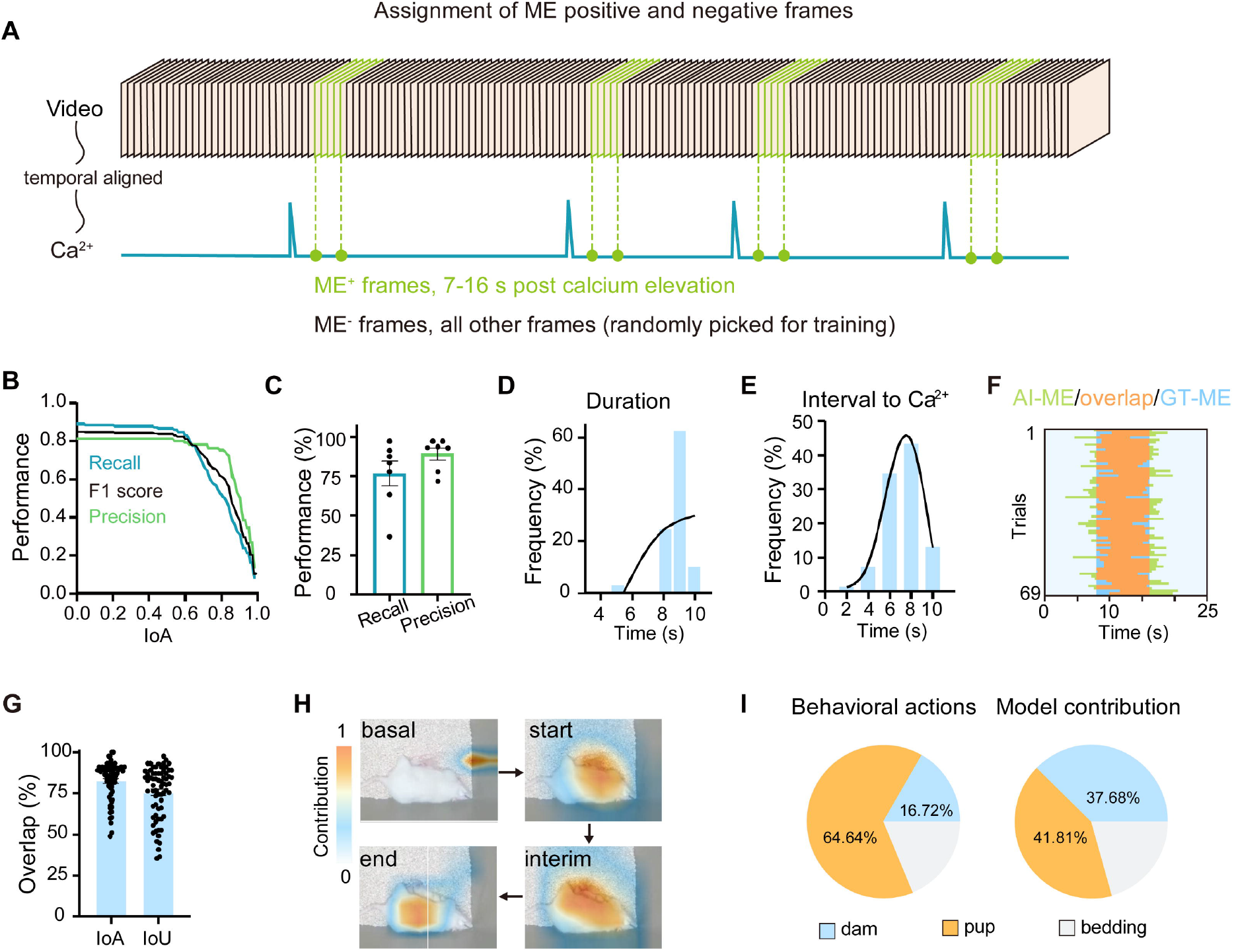
Dam-pup interactions contribute to the recognition by ME Decoder. **(A)**Ilustration of the selection of positive and negative training inputs. **(B)** The performance of ME Decoder in the validation dataset after final training (n = 6 videos). **(C)** Performance of the ME Decoder in test dataset (n = 7 videos). **(D)** Histograms showing the duration of Ai-MEs in test dataset. **(E)** Histograms showing the intervals between the onset of oxytocin calcium waves and Ai-ME in test dataset. **(F)** Temporal overlap heatmap between Ai-ME and GT-ME in test dataset. Green, Ai-ME exclusive; orange, Ai-ME overlapped with ADPI; blue, GT-ME exclusive. **(G)** Percentage of the overlap between Ai-ME and GT-ME. IoA, Intersection over area (Overlap duration) / (Ai-ME duration); IoU, intersection over union, (Overlap duration) / (Total duration of Ai-ME and GT-ME). **(H)** Regional contribution to ME Decoder’s decision visualized by heatmaps. **(I)** Percentage of behavioral actions (left, frame differences) and discriminative contributions (right, weights in model) in ME Decoder’s decision for dam (blue), pup (orange), and bedding (gray) (n = 7 videos).

To understand the features that ME Decoder learned through training, we used Grad-CAM++ to generate spatiotemporal saliency maps and visualize the model’s attention during classification [51]. These maps highlighted the dam’s back and the interactive interfaces between the dam and pups, including her abdomen and all pups, suggesting these regions are critical for the model’s decision (Fig 4H). By further segmenting video frames into three regions: dam, pups, and bedding, we found that the dam (37.7 ± 0.3%) and pups (41.8 ± 0.5%) exhibited comparable contributions to the model’s decision, despite the limited mobility of the dam (16.7 ± 0.6%; Fig 4I). These results underscore the importance of integrating cues from the dam and pups to recognize milk ejections in conscious animals.

### Interpretable behavioral signatures define milk ejection

ME Decoder highlights dam-pup interacting areas as salient features of milk ejection. To gain further insights to the specific behavioral changes and movements associated with ME, we next performed fine-grained analyses of behavioral dynamics and divided behavioral changes occurring during lactation into three different categories for either dams (D1-D3) or pups (P1-P3) based on the head and back movement of the dam, as well as that of pups’ bodies and limbs (Fig 5A–5B, Video S1-S6, detailed in methods). To emphasize the dynamic nature of these events, the basal quiescent state of lactation, with no apparent movement of both dam and pups was referred to D0 or P0 (Video S7).

**Fig 5.**
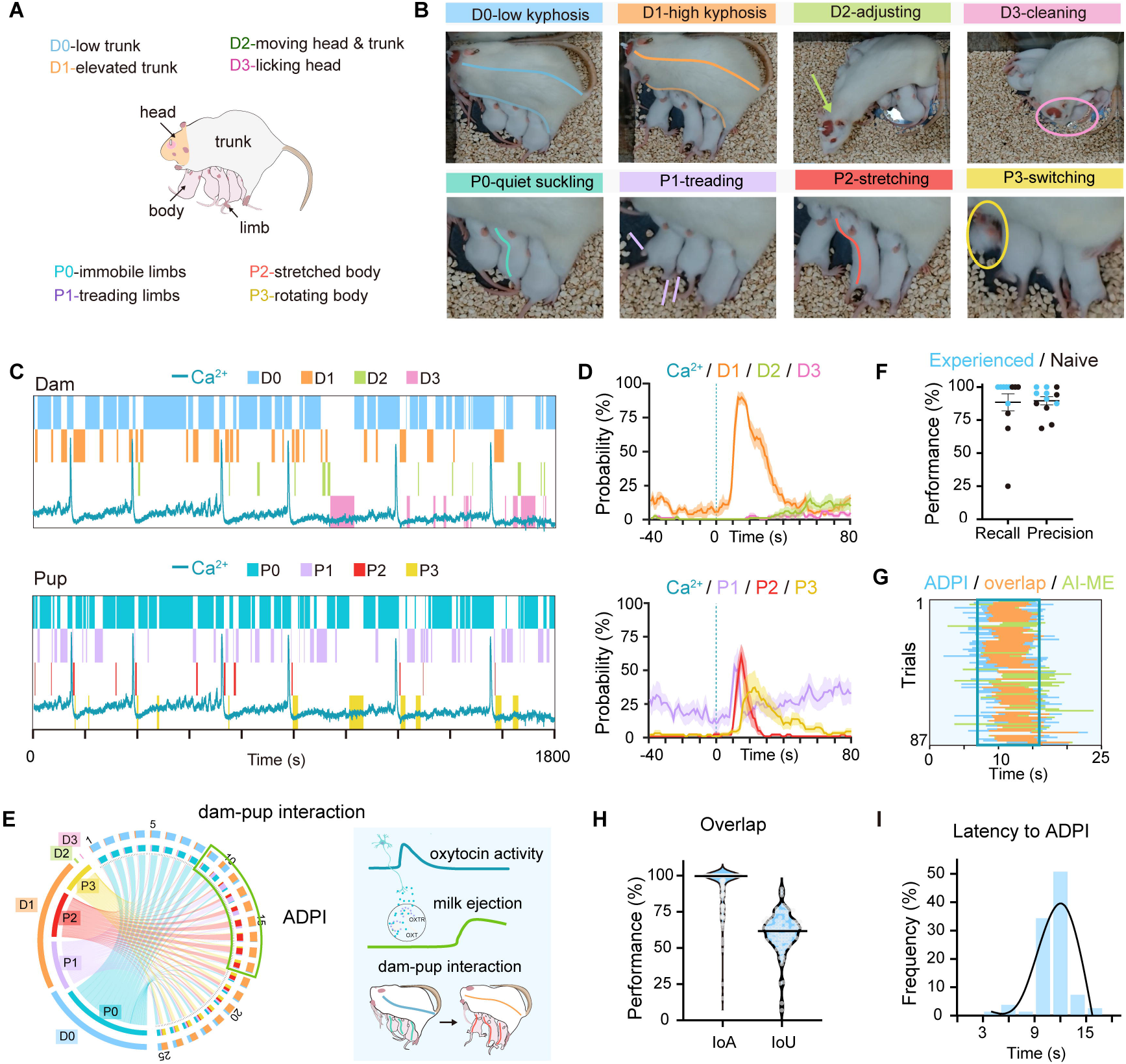
Interpretable behavioral signatures define milk ejection. **(A-B)** Schematic illustration (A) and representative frames (B) showing different of behavioral categories of dam and pups during suckling. **(C)** Representative traces showing the calcium waves of oxytocin neurons (blue line), behavioral types of the dam (top) and pup (bottom) during 30 min suckling. **(D)** Mean probability of different behaviors in dam (top) and pups (bottom) aligned with the timing of the onset of oxytocin calcium waves (n = 6 videos, 77 trials). **(E)** Left, Circos plots displaying behavioral composition during ME episodes, with behavioral categories shown on the left, whereas timing relative to oxytocin neuron calcium activity on the right. Right, Schematic summary demonstrating the relationship of oxytocin neuronal activity, milk ejection and ADPI. **(F)** Recall and precision rates for ME recognition using ADPI by both experienced and naive experimenters. (n = 6 videos, 87 trials). **(G)** Heatmap showing the overlapped time regions between ADPI, Ai-ME and GT-ME. Blue, ADPI-exclusive; orange, ADPI overlapped with GT-ME; green, Ai-ME-exclusive; the blue box indicate the GT-ME. **(H)** Percentage of the overlap between ADPI and GT-ME. IoA, Intersection over area, (Overlap duration) / ADPI duration); IoU, intersection over union, (Overlap duration) / (Total duration of GT-ME and ADPI). **(I)** Histograms showing the latency of ADPI to onset of calcium wave.

We then constructed an ethogram by temporally aligning these categories with oxytocinergic calcium signals. In a short time period after calcium waves when milk ejections were anticipated (8-17 s), the dam and pups constantly exhibited highly coordinated behavioral interactions, starting from pronounced high kyphosis of the dam (D1), followed by pup treading (P1) and stretching (P2), and typically ending with pup switching (P3) (sequence: D1-P1-P2-P3) (Fig 5C-E and Fig S2A-B). Such behavioral sequences were exclusively associated with positive training video segments and absent in negative segments (Fig S2C-D).

Given its tight coupling with oxytocinergic activity, we referred to this behavioral pattern as Activity-coupled Dam-Pup Interactions (ADPI) (Fig 5E and Video S8). Notably, the regions where ADPI occurred were consistent with the attention regions on which ME Decoder focused (Fig 4H), as generated by the Class Activation Mapping (CAM) to demonstrate the ME Decoder focus. The spatial correlation indicates that ADPI represents the interpretable and discriminative features learned by the model for classification.

Next, to assess whether ADPI could serve as a reliable behavioral proxy for milk ejection, two investigators (one experienced and the other naïve) independently assessed milk ejections from videos using ADPI. Concurrently recorded calcium signals of oxytocin neurons were blinded to the investigators and served as the ground truth. Both investigators achieved high recall (88.4 ± 22.5%) and precision (89.6 ± 10.5%) for the assessments (Fig 5F). Moreover, milk ejections identified by ADPI closely overlapped with ground truth and automatically identified (Ai)-ME (Fig 5G-H), and occurred with a consistent latency from the onset of calcium waves (11.1 ± 0.2 s, Fig 5I). These results establish ADPI as a reliable, observer-independent behavioral signature for milk ejection recognition and provides interpretability for the decision of the ME decoder.

### Behavioral modeling detects changes of milk ejections

Peripheral action of oxytocin on mammary myoepithelial cells is required for milk ejections [6]. To determine the accuracy of ME-Decoder and ADPI in ME measurement in different condition, we intraperitoneally administered the oxytocin receptor antagonist, L-368,899 (5 mg/kg) to lactating rats (Fig 6A), which is known to inhibit milk ejections [52], verified by the significantly reduced pup weight gain (Fig 6B). Notably, both manual analyses of ADPI and automatic ME Decoder successfully detected the reduced numbers of milk ejections (Fig 6C-6E), suggesting that they were sensitive in detecting changes in milk ejections under different conditions (Fig 6F). Because L-368,899 has a high blood brain barrier penetrance [53], we further tested its effects on oxytocinergic neuronal activity in OXT-Cre dams. While milk ejections were reduced (Fig 6G-I), oxytocin neurons remained unchanged, with unaffected interval, amplitude, or duration of calcium waves (Fig 6J-O). These data indicate that L-368,899 inhibited milk ejection without compromising central oxytocinergic neuronal activity.

**Fig 6.**
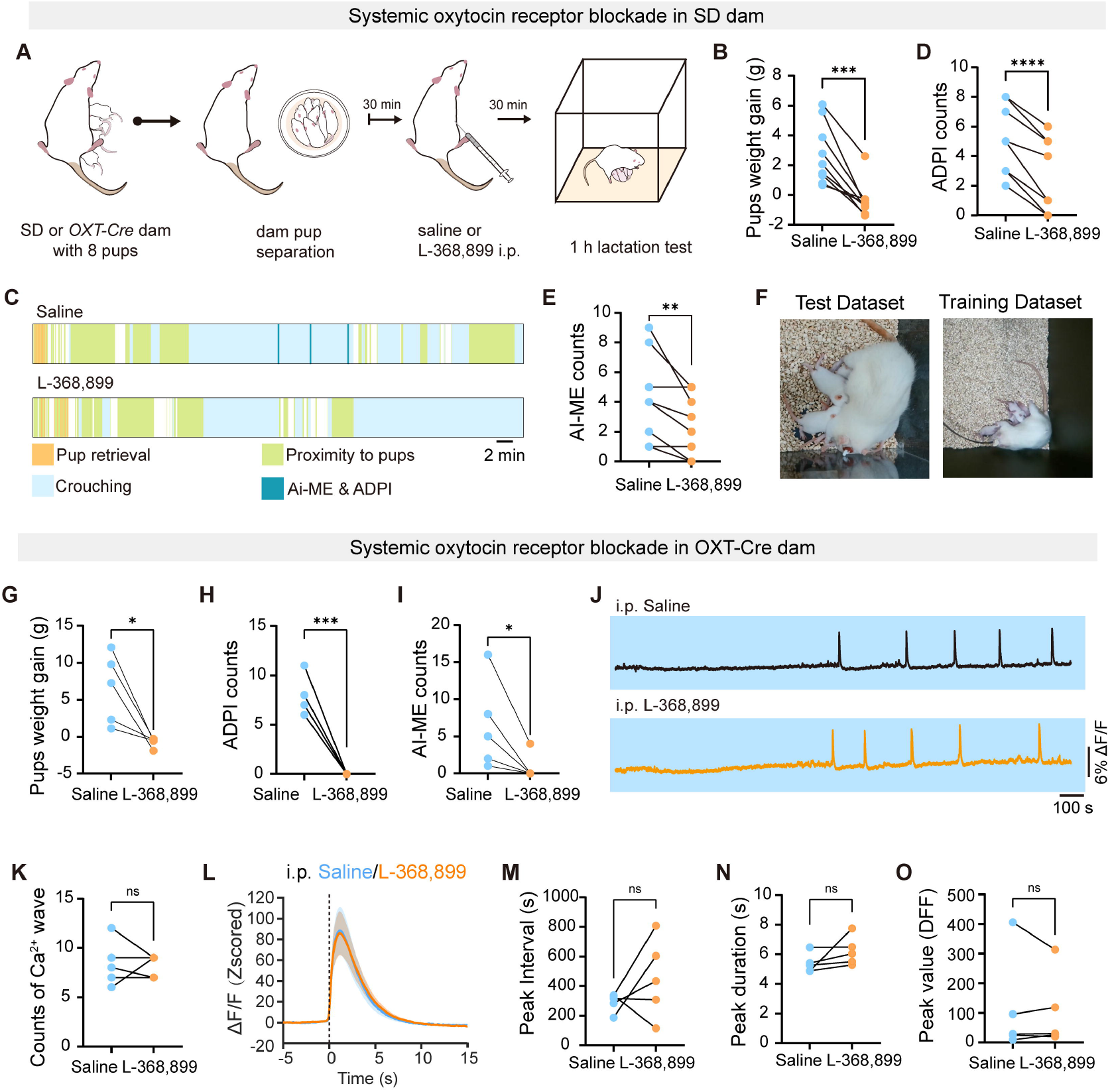
Behavioral modeling detects changes of milk ejections. **(A)** Schematic illustration of procedures of systemic oxytocin receptor blockade. **(B)** Pup weight gain of dam crouching over pups during 1 h lactation in the absence (saline) or presence of oxytocin receptor antagonist (L-368,899) (n = 9 litters). *** *p* < 0.001 by two-tailed paired t-test. **(C)** Ethograms of dam behavior before and after systematic oxytocin receptor blockade. **(D-E)** Number of manual annotated ADPIs (D) and ME Decoder identified Ai-ME (E) during 1 h lactation (saline versus L-368,899, n = 9 litters). **** *p* < 0.0001, ** *p* < 0.01 by two-tailed paired t-test. **(F)** Video frames from the training dataset and the test dataset. **(G-I)** Number of pup weight gain (G), ADPI (H), and Ai-ME (I) during 1 h lactation, before and after systematic oxytocin receptor blockade, tested in OXT-Cre dams (n = 5 dams). * *p* < 0.05, *** *p* < 0.001 by two-tailed paired t-test **(J)** Representative calcium traces of oxytocin neurons during lactation, before and after systematic oxytocin receptor blockade in OXT-Cre dam. **(K)** Quantification of calcium number before and after systematic oxytocin receptor blockade (n = 5 dams). *p* = 0.8466 by two-tailed paired t-test **(L)** Average (mean ± SEM) calcium wave of oxytocin neurons during suckling, before and after systematic oxytocin receptor blockade. **(M-O)** Quantification of the interval (M), duration (N), and height (O) of oxytocin neurons during lactation, before and after systematic oxytocin receptor blockade (n = 5 dams). *p* = 0.7794 for M, *p* = 0.1829 for N, *p* = 0.6217 for O by two-tailed paired t-test

## Discussion

Understanding lactation is important for reproductive health. In the present study, we developed a supervised deep-learning framework (ME Decoder) to allow automated and noninvasive detection of milk ejections, providing a novel approach for quantitative measurements of lactation. ME Decoder learned from the sequential behavioral responses of ADPI, manifested by high kyphosis of the dam, followed by pup treading and stretching, for discriminative localization of milk ejections. Using these new tools, we verified that systematically blocking oxytocin signaling inhibits milk ejections but do not disturb milk ejection-triggering pulsatile oxytocinergic firing. These findings provide important insights into mechanistic understanding the interplay between oxytocin neuron activity and behavioral output of natural lactation in freely behaving rats and presents a generalizable framework for investigating lactation-related physiological and emotional impacts.

### Automation of milk ejection analyses

A comprehensive understanding of lactation requires a methodological paradigm that is both precise and applicable to natural conditions. Traditional approaches, IMP recording provides direct physiological readout of milk ejection but necessitates anesthesia [52], which precludes faithful representation of the full suite of physiological processes in conscious animals [54]. Machine learning has emerged as a powerful approach for automating behavioral analysis [20]. However, due to multi-animal interactions and frequent visual occlusion of pups, attempts using skeleton-based approaches (e.g. DeepLabcut) were unsuccessful [22, 55]. Therefore, we developed a new strategy to monitor and quantify behavior during suckling, ME Decoder.

ME Decoder is a computer vision-based deep learning model that combines architectures for feature detection and refinement, enabling dynamic and complex behavioral analyses involving multiple animals. This framework is user-friendly and cost-effective, requiring only overhead-view videos as input. The adaptability of this framework has been validated in different experimental conditions, including variations in environmental settings and animal models, establishing ME Decoder as a versatile resource to investigate lactation. Training the framework with different datasets will expand its application in additional ethological analyses other than lactation.

### Behavioral signatures of lactation

Breastfeeding is a universal feature of all mammals and constitutes a critical component of maternal behavior. While other maternal actions such as pup retrieval have been extensively studied [40, 56, 57], much less attention has been paid to behavioral features of lactation. Because behavioral manifestations of milk ejection remain unclear, quantitative and systematic ethological analyses of lactation are challenging. While pup stretch has been suggested to indicate milk ejection [49], discriminating rigid stretch form other pup movements is challenging during natural lactation, as its occurrence, representation and duration varies highly. By contrast, the high kyphosis (D1) of the dam and switching of pups (P3) are readily recognized. We noticed that when using D1 and P3 as references, it becomes much easier to recognize pup treading (P1) and stretching (P2) often occurs in-between these two actions.

Inspired by the spatiotemporal features learned by the ME Decoder, we further defined these interpretable behavioral sequences (P1-D1-D2-D3), namely, ADPI, as representations of milk ejections. ADPI, which integrates characteristic actions from both the mother and offspring, transforms the ambiguous individual actions into a robust behavioral pattern. This enables precise analyses of milk ejections and reduces the needs for invasive IMP and neuronal recordings, and serves as an important complementation and verification for ME Decoder analyses.

### Neural modulation of milk ejection

The unique burst firing of oxytocin neurons during suckling has been a subject of intense interest. Central oxytocin signaling has been proposed to facilitate burst firing of oxytocin neurons via either synaptic or extra synaptic transmission [58–61]. These studies were mainly conducted in lactating rats under anesthesia, a brain state with globally inhibited neuronal activity. With new approaches developed in the present study, we demonstrated that systematically blocking oxytocin signaling had little effects on the burst firing of oxytocin neurons. While these data need to be further investigated using genetic ablation of oxytocin receptors in neurons, our observation is consistent with previous reports that systematic injection of another oxytocin receptor antagonist (F-792) did not affect the burst firing of oxytocin neurons [62]. However, both intra-cerebroventricular and intra-SON infusion of oxytocin receptor antagonist appeared to reduce plasma oxytocin and milk transfer [60, 63]. These data indicate that central oxytocinergic activity during milk ejection may be regulated at multiple levels. It is likely that the burst firing of oxytocin neurons is regulated by upstream inputs [13, 64, 65], which initiate and coordinate population-wide synchrony of oxytocin neurons during lactation.

### Limitations of the study

Fiber photometry recording detects the calcium signals at population level, thus may compromise the detection sensitivity and specificity. In addition, fiber location and virus expression may also increase the variation of recorded signals. Miniscope-based single-cell *in vivo* imaging or direct multi-single unit recording of action potential firing are alternative strategies that detect neuronal activity at single cell level to enable precise investigation of neuron-to neuron synchronization within burst. Moreover, the datasets used for training ME Decoder were primarily from the middle and late stages of lactation (PPD7-PPD20). Further optimization may be required for its application to earlier lactation phases, e.g. from PPD1-PPD6.

## ACKNOWLEDGEMENTS

We thank Dr. Shan Jiang and Dr. Enyin Lai at Zhejiang University for assistance in IMP recording, Dr. Yufeng Wang at Zhejiang Chinese Medical University for suggestions on lactation related recording.

This work was supported by the National Science and Technology Innovation 2030-Major Projects (2021ZD0202700 to Z.G.), National Natural Science Foundation of China (82288101 to S.D.), National Science and Technology Innovation 2030-Major Projects (2021ZD0200400 to G.P.), National Natural Science Foundation of China (82090033 to S.D., 32371003 and 32070974 to Z.G.), Key R&D Program of Zhejiang Province (2024SSYS0017 to S.D.), National High-Level Talent Special Support Programs (10,000 Talents Program) Leading Talents to Z.G, and National Natural Science Foundation of China (61925603 to G.P., 62376247 to Q.Z.).

## Author contributions

Conceptualization, Z.G., W.X., Q.Z, S.D.; Methodology, W.X.; Model construction, Y.W., W.X., Q.Z., G.P.; Data Analysis, W.X., Y.Y., B.S., Y.G., Y.C., T.Z.; Validation, Y.G., T.Z.; Investigation, W.X.; Writing – Original Draft, W.X., Z.G., Y.W.; Writing – Review & Editing, Z.G., W.X., C.B., Q.Z., B.Z., L.Q.; Visualization, W.X., Y.W.; Supervision, Z.G., S.D., Q.Z. G.P.; Project Administration, Z.G., W.X.; Funding Acquisition, Z.G., S.D., Q.Z., G.P..

## Competing interests

Authors declare that they have no competing interests.

## Data and materials availability

All data needed to evaluate the conclusions in the paper are present in the paper and/or the Supplementary Materials. All data reported in this paper will be shared by the lead contact upon request. The codes used in this study are available at GitHub (https://anonymous.4open.science/r/MEDecoder) with a LGPL-3.0 License.

---

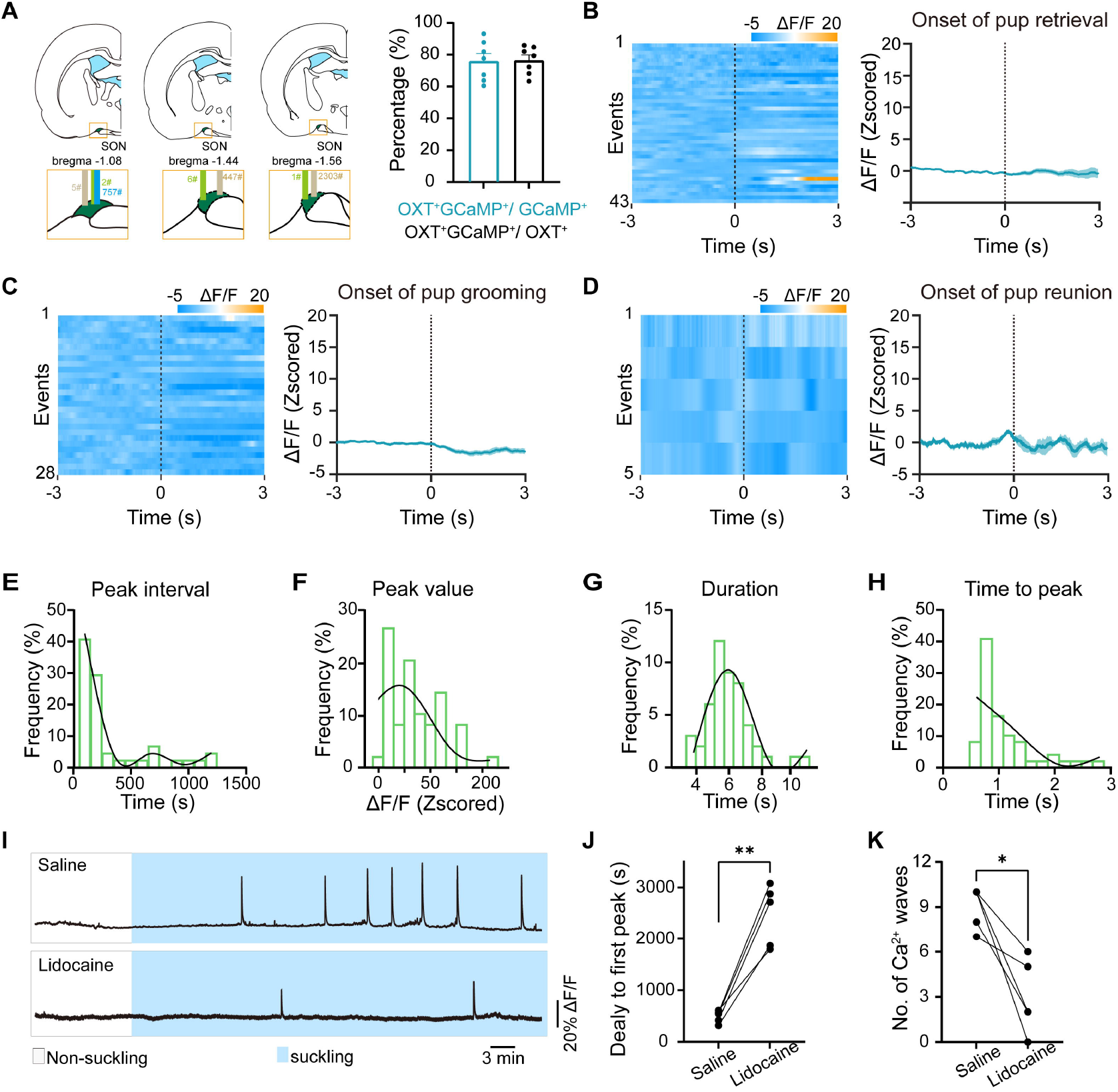

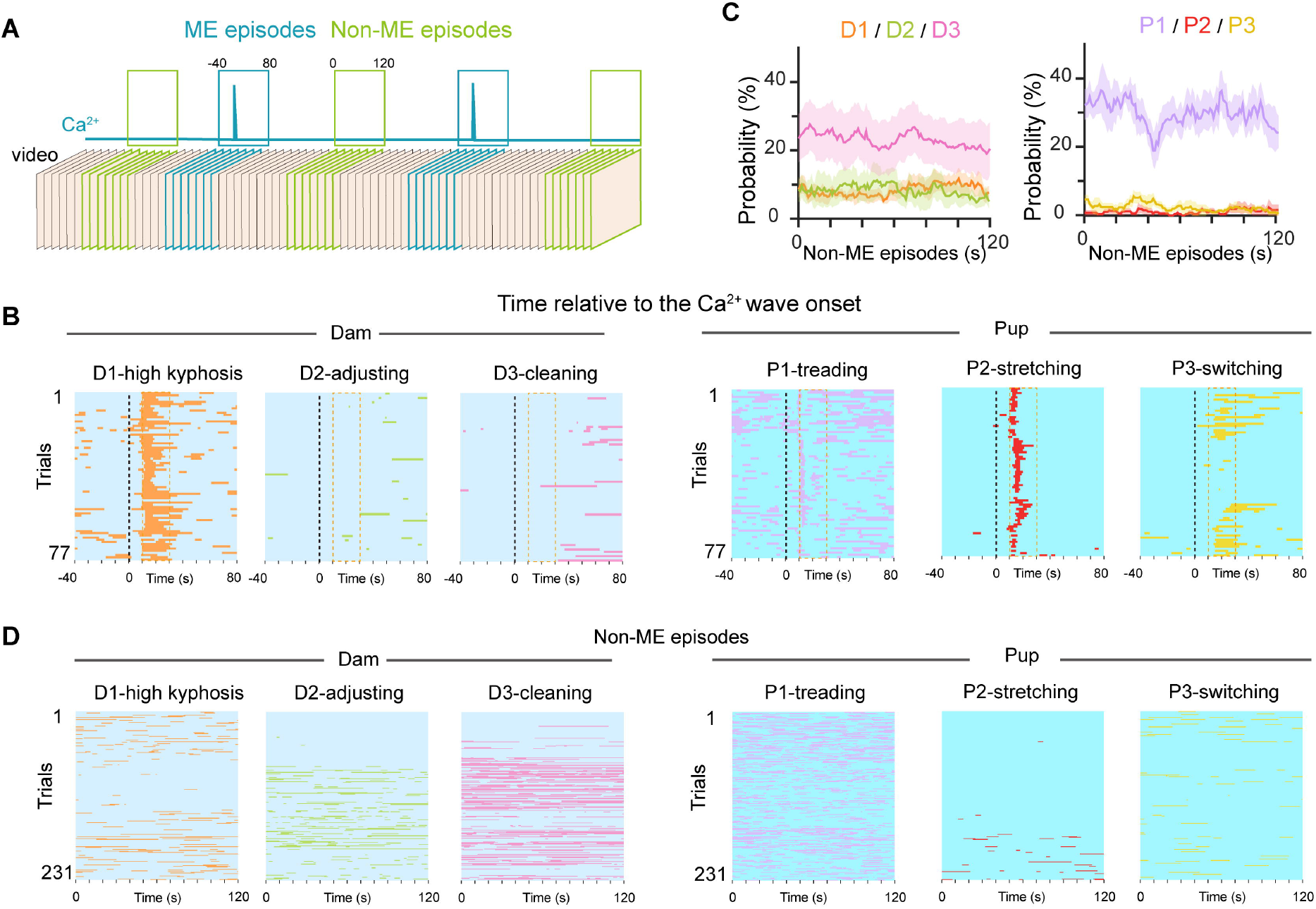

## Notes

### Competing Interest Statement

The authors have declared no competing interest.

## REFERENCES

[1] Gaylord SG. The principles of classification and a classfication of mammals. New York: American Museum of Natural History, 1945.

[2] Harris GW. The central nervous system, neurohypophysis and milk ejection. Proc R Soc Lond B Biol Sci 1958, 149: 336–353.

[3] Lincoln DW, Paisley AC. Neuroendocrine control of milk ejection. Reproduction 1982, 65: 571–586.

[4] Wakerley JB, Lincoln DW. Intermittent release of oxytocin during suckling in the rat. Nat New Biol 1971, 233: 180–181.

[5] Neville MC. Anatomy and physiology of lactation. Pediatr Clin N Am 2001, 48: 13–34.

[6] Soloff MS. Oxytocin receptors and mammary myoepithelial cells. J Dairy Sci 1982, 65: 326–337.

[7] Wakerley JB, Lincoln DW. The milk-ejection reflex of the rat: A 20- to 40-fold acceleration in the firing of paraventricular neurones during oxytocin release. J Endocrinol 1973, 57: 477–493.

[8] Summerlee AJS, Lincoln DW. Electrophysiological recordings from oxytocinergic neurones during suckling in the unanaesthetized lactating rat. J Endocrinol 1981, 90: 255–265.

[9] Belin V, Moos F, Richard P. Synchronization of oxytocin cells in the hypothalamic paraventricular and supraoptic nuclei in suckled rats: Direct proof with paired extracellular recordings. Exp Brain Res 1984, 57: 201–203.

[10] Belin V, Moos F. Paired recordings from supraoptic and paraventricular oxytocin cells in suckled rats: Recruitment and synchronization. J Physiol 1986, 377: 369–390.

[11] Armstrong WE. Central nervous system control of oxytocin secretion during lactation. Knobil and Neill’s Physiology of Reproduction. Amsterdam: Elsevier, 2015: 527–563.

[12] Lincoln DW, Wakerley JB. Electrophysiological evidence for the activation of supraoptic neurones during the release of oxytocin. J Physiol 1974, 242: 533–554.

[13] Yukinaga H, Hagihara M, Tsujimoto K, Chiang HL, Kato S, Kobayashi K, et al. Recording and manipulation of the maternal oxytocin neural activities in mice. Curr Biol 2022, 32: 3821–3829.e6.

[14] Yaguchi K, Miyamichi K, Tasaka GI. Flexible adjustment of oxytocin neuron activity in mouse dams revealed by microendoscopy. Sci Adv 2024, 10: eadt1555.

[15] Thirtamara Rajamani K, Leithead AB, Kim M, Barbier M, Peruggia M, Niblo K, et al. Efficiency of cell-type specific and generic promoters in transducing oxytocin neurons and monitoring their neural activity during lactation. Sci Rep 2021, 11: 22541.

[16] Perkinson MR, Kim JS, Iremonger KJ, Brown CH. Visualising oxytocin neurone activity in vivo: The key to unlocking central regulation of parturition and lactation. J Neuroendocrinol 2021, 33: e13012.

[17] Yaguchi K, Hagihara M, Konno A, Hirai H, Yukinaga H, Miyamichi K. Dynamic modulation of pulsatile activities of oxytocin neurons in lactating wild-type mice. PLoS One 2023, 18: e0285589.

[18] Krol KM, Grossmann T. Psychological effects of breastfeeding on children and mothers. Bundesgesundheitsblatt Gesundheitsforschung Gesundheitsschutz 2018, 61: 977–985.

[19] Kim P, Feldman R, Mayes LC, Eicher V, Thompson N, Leckman JF, et al. Breastfeeding, brain activation to own infant cry, and maternal sensitivity. J Child Psychol Psychiatry 2011, 52: 907–915.

[20] Anderson DJ, Perona P. Toward a science of computational ethology. Neuron 2014, 84: 18–31.

[21] Gomez-Marin A, Paton JJ, Kampff AR, Costa RM, Mainen ZF. Big behavioral data: Psychology, ethology and the foundations of neuroscience. Nat Neurosci 2014, 17: 1455–1462.

[22] Nath T, Mathis A, Chen AC, Patel A, Bethge M, Mathis MW. Using DeepLabCut for 3D markerless pose estimation across species and behaviors. Nat Protoc 2019, 14: 2152–2176.

[23] Hu K, Jin J, Zheng F, Weng L, Ding Y. Overview of behavior recognition based on deep learning. Artif Intell Rev 2023, 56: 1833–1865.

[24] Mathis MW, Mathis A. Deep learning tools for the measurement of animal behavior in neuroscience. Curr Opin Neurobiol 2020, 60: 1–11.

[25] Mathis A, Mamidanna P, Cury KM, Abe T, Murthy VN, Mathis MW, et al. DeepLabCut: Markerless pose estimation of user-defined body parts with deep learning. Nat Neurosci 2018, 21: 1281–1289.

[26] Pereira TD, Tabris N, Matsliah A, Turner DM, Li J, Ravindranath S, et al. SLEAP: A deep learning system for multi-animal pose tracking. Nat Meth 2022, 19: 486–495.

[27] Bohnslav JP, Wimalasena NK, Clausing KJ, Dai YY, Yarmolinsky DA, Cruz T, et al. DeepEthogram, a machine learning pipeline for supervised behavior classification from raw pixels. Elife 2021, 10: e63377.

[28] Serre T. Deep learning: The good, the bad, and the ugly. Annu Rev Vis Sci 2019, 5: 399–426.

[29] Zhang B, Qiu L, Xiao W, Ni H, Chen L, Wang F, et al. Reconstruction of the hypothalamo-neurohypophysial system and functional dissection of magnocellular oxytocin neurons in the brain. Neuron 2021, 109: 331–346.e7.

[30] Grosvenor CE, Turner CW. Estimation of amount of oxytocin released as result of nursing stimuli in lactating rat. Exp Biol Med 1957, 95: 131–133.

[31] Vorherr H, Kleeman CR, Lehman E. Oxytocin-induced stretch reaction in suckling mice and rats: A semiquantitative bio-assay for oxytocin. Endocrinology 1967, 81: 711–715.

[32] Peng W, Wu Z, Song K, Zhang S, Li Y, Xu M. Regulation of sleep homeostasis mediator adenosine by basal forebrain glutamatergic neurons. Science 2020, 369: eabb0556.

[33] Peng W, Liu X, Ma G, Wu Z, Wang Z, Fei X, et al. Adenosine-independent regulation of the sleep –wake cycle by astrocyte activity. Cell Discov 2023, 9: 16.

[34] Zhang C, Zhu H, Ni Z, Xin Q, Zhou T, Wu R, et al. Dynamics of a disinhibitory prefrontal microcircuit in controlling social competition. Neuron 2022, 110: 516–531.e6.

[35] Osakada T, Yan R, Jiang Y, Wei D, Tabuchi R, Dai B, et al. A dedicated hypothalamic oxytocin circuit controls aversive social learning. Nature 2024, 626: 347–356.

[36] Lincoln DW, Hill A, Wakerley JB. The milk-ejection reflex of the rat: An intermittent function not abolished by surgical levels of anaesthesia. J Endocrinol 1973, 57: 459–476.

[37] Xia H, Zhan Y. A survey on temporal action localization. IEEE Access 2020, 8: 70477–70487.

[38] Lin J, Gan C, Wang K, Han S. TSM: Temporal shift module for efficient and scalable video understanding on edge devices. IEEE Trans Pattern Anal Mach Intell 2022, 44: 2760–2774.

[39] Lin T, Liu X, Li X, Ding E, Wen S. BMN: Boundary-matching network for temporal action proposal generation. 2019 IEEE/CVF International Conference on Computer Vision (ICCV). October 27-November 2, 2019. Seoul, Korea. IEEE, 2019: 3888–3897.

[40] Marlin BJ, Mitre M, D’amour JA, Chao MV, Froemke RC. Oxytocin enables maternal behaviour by balancing cortical inhibition. Nature 2015, 520: 499–504.

[41] Carcea I, Caraballo NL, Marlin BJ, Ooyama R, Riceberg JS, Mendoza Navarro JM, et al. Oxytocin neurons enable social transmission of maternal behaviour. Nature 2021, 596: 553–557.

[42] Valtcheva S, Issa HA, Bair-Marshall CJ, Martin KA, Jung K, Zhang Y, et al. Neural circuitry for maternal oxytocin release induced by infant cries. Nature 2023, 621: 788–795.

[43] Yuan Y, Gao Z, Xiao W. The role of oxytocin in parental care. Endocrinology 2025, 166: bqaf129.

[44] Tang Y, Benusiglio D, Lefevre A, Hilfiger L, Althammer F, Bludau A, et al. Social touch promotes interfemale communication via activation of parvocellular oxytocin neurons. Nat Neurosci 2020, 23: 1125–1137.

[45] Gawali VS, Lukacs P, Cervenka R, Koenig X, Rubi L, Hilber K, et al. Mechanism of modification, by lidocaine, of fast and slow recovery from inactivation of voltage-gated Na+ channels. Mol Pharmacol 2015, 88: 866–879.

[46] Ueda T, Yokoyama Y, Irahara M, Aono T. Influence of psychological stress on suckling-induced pulsatile oxytocin release. Obstet Gynecol 1994, 84: 259–262.

[47] Del Ciampo IRL, Del Ciampo LA. Breastfeeding and the benefits of lactation for women’s health. Rev Bras Ginecol Obstet 2018, 40: 354–359.

[48] Robinson SR, Smotherman WP. Organization of the stretch response to milk in the rat fetus. Dev Psychobiol 1992, 25: 33–49.

[49] Drewett RF, Statham C, Wakerley JB. A quantitative analysis of the feeding behaviour of suckling rats. Anim Behav 1974, 22: 907–913.

[50] Lin T, Liu X, Li X, Ding E, Wen S. BMN: Boundary-matching network for temporal action proposal generation. 2019: arXiv: 1907.09702. https://arxiv.org/abs/1907.09702

[51] Chattopadhay A, Sarkar A, Howlader P, Balasubramanian VN. Grad-CAM++: Generalized gradient-based visual explanations for deep convolutional networks. 2018 IEEE Winter Conference on Applications of Computer Vision (WACV). March 12–15, 2018, Lake Tahoe, NV, USA. IEEE, 2018: 839 –847.

[52] Wakerley JB, Dyball REJ, Lincoln DW. Milk ejection in the rat: The result of a selective release of oxytocin. J Endocrinol 1973, 57: 557–558.

[53] Freeman SM, Catrow JL, Cox JE, Turano A, Rich MA, Ihrig HP, et al. Binding affinity, selectivity, and pharmacokinetics of the oxytocin receptor antagonist L-368, 899 in the coyote (Canis latrans). Comp Med 2024, 74: 3–11.

[54] Eisen AJ, Kozachkov L, Bastos AM, Donoghue JA, Mahnke MK, Brincat SL, et al. Propofol anesthesia destabilizes neural dynamics across cortex. Neuron 2024, 112: 2799–2813.e9.

[55] Hsu AI, Yttri EA. B-SOiD, an open-source unsupervised algorithm for identification and fast prediction of behaviors. Nat Commun 2021, 12: 5188.

[56] Schiavo JK, Valtcheva S, Bair-Marshall CJ, Song SC, Martin KA, Froemke RC. Innate and plastic mechanisms for maternal behaviour in auditory cortex. Nature 2020, 587: 426–431.

[57] Cui X, Xiao L. Complexity of the hypothalamic oxytocin system and its involvement in brain functions and diseases. Neurosci Bull 2025, 41: 1267–1288.

[58] Lambert RC, Dayanithi G, Moos FC, Richard P. A rise in the intracellular Ca2+ concentration of isolated rat supraoptic cells in response to oxytocin. J Physiol 1994, 478: 275–287.

[59] Li D, Liu H, Liu X, Wang H, Li T, Wang X, et al. Involvement of hyperpolarization-activated cyclic nucleotide-gated channel 3 in oxytocin neuronal activity in lactating rats with pup deprivation. ASN Neuro 2020, 12: 1759091420944658.

[60] Freund-Mercier MJ, Richard P. Electrophysiological evidence for facilitatory control of oxytocin neurones by oxytocin during suckling in the rat. J Physiol 1984, 352: 447–466.

[61] Freund-mercier MJ, Moos F, Poulain DA, Richard P, Rodriguez F, Theodosis DT, et al. Role of central oxytocin in the control of the milk ejection reflex. Brain Res Bull 1988, 20: 737–741.

[62] Leng G, Russell JA. The peptide oxytocin antagonist F-792, when given systemically, does not act centrally in lactating rats. J Neuroendocrinol 2016, 28: 12331.

[63] Neumann I, Koehler E, Landgraf R, Summy-Long J. An oxytocin receptor antagonist infused into the supraoptic nucleus attenuates intranuclear and peripheral release of oxytocin during suckling in conscious rats. Endocrinology 1994, 134: 141–148.

[64] Ingrarn CD, Adams TST, Jiang QB, Terenzi MG, Lambert RC, Wakerley JB, et al. Limbic regions mediating central actions of oxytocin on the milk-ejection reflex in the rat. J Neuroendocrinol 1995, 7: 1 –13.

[65] Meddle SL, Leng G, Selvarajah JR, Bicknell RJ, Russell JA. Direct pathways to the supraoptic nucleus from the brainstem and the main olfactory bulb are activated at parturition in the rat. Neuroscience 2000, 101: 1013–1021.

