## supporting information for "Decoding the oxytocinergic and behavioral signatures of milk ejection"

**Fig S1. Oxytocin neurons showed no apparent signal during maternal behaviors. (A)** Fiber location (left) and co-labeling of oxytocin and GCaMP6m in the SON. 76.10% of the GCaMP6m-labeled cells were positive for OXT-NP, with 76.64% of OXT positive cells labeled by GCaMP6m (n = 7 rats; 561 cells). **(B-D)** Individual (left, heatmap) and average (right, mean ± SEM) calcium signals aligned to the onset of pup retrieval (B), pup grooming (C) and pup reunion (D). **(E-H).** Frequency distribution of calcium waves during suckling for inter-peak intervals (E), peak amplitude (F), peak duration (G), and time to peak (H) (n = 5 dams, 49 trials). **(I)** Representative traces of oxytocin calcium signals during suckling after local anesthesia by subcutaneous injection of lidocaine to all nipples. **(J-K)** Prolonged delay to first peak (J) after suckling and reduced numbers of calcium waves (K) after local anesthesia of the nipples.

**Fig S2. Behavioral characterization of dam and pups during suckling. (A)** Schematic illustrations of defining ME and Non-ME episodes using calcium waves as a reference. ME episodes were defined as episodes of 40 s pre- and 80 s post-calcium waves, whereas Non-ME episodes were segments of 120 s from the remaining videos. **(B)** Heatmaps showing the temporal distribution of individual behavioral categories aligned to the onset of oxytocinergic calcium waves in ME episodes (n = 6 videos, 77 trials). **(C)** Mean probability of different behaviors in the dam (left) and pups (right) in Non-ME events (n = 6 videos, 231 trials). **(D)** Heatmaps showing the temporal distribution of individual behavioral categories in Non-ME episodes (n = 6 videos, 231 trials).

**Video S1. dam high kyphosis.**

**Video S2. dam adjusting.**

**Video S3. dam cleaning.**

**Video S4. pup treading.**

**Video S5. pup stretching.**

**Video S6. pup switching.**

**Video S7. basal states of dam and pups.**

**Video S8. ADPI**

**Methods**

**Methods of ME Decoder** **Training**

The TSM is composed of a ResNet50 backbone and a single-layer fully connected (FC) classification network [1]. The backbone network generates behavioral representations for each frame and the FC network determines a segment (the sequence of behavioral representation, defined by start and end timestamps) as ME or not. The BMN, which is implemented based on multi-layer 3D convolutions [2], locates candidate ME segments.

**Data Preprocessing.** Because dam-pup interactions are gradual across consecutive frames, a lower frame rate is sufficient for behavior analysis. Thus, we reduced the frame rate of collected videos to 5 frames per second (FPS) using FFmpeg (https://ffmpeg.org). Also, because the dam and pups occupy only a small portion (~1/6) of the video frame, we used a deep learning tool, PaddleDetection (https://github.com/PaddlePaddle/PaddleDetection), to automatically crop a rectangular region centered around the dam. This region covers 1/4 of the frame (half of its width and height, with a final resolution of 320x240 pixels) to ensure the dam and pups are included.

**Data Partition.** We collected 63 videos containing calcium recording and split them into three subsets: 50 for training, 6 for validation, and 7 for testing. The training dataset was used to train deep neural networks, the validation dataset was used to find the optimal network parameters, and the test dataset was used to assess network performance. Video source details are provided in the supplementary table 1.

**Data Annotation.** We annotated the start and end timestamps of ME segments for each collected video using recorded calcium signals as reference (7-16 s post calcium wave onset). All frames within each segment are automatically labelled as milk ejection frames, while the remaining frames are labelled as non-milk ejection frames. Following the method [3], we generated two probability vectors indicating frame-level probabilities of being ME boundaries, and a probability matrix indicating segment-level ME probability. Specifically, the probability vectors represent each frame's likelihood of being ME's start/end timestamp and each element of the probability matrix represents each segment's likelihood of being ME, where row index and column index denote the start/end timestamp, respectively. In total, we obtained 754 ME segments in the training subset, 103 in the validation subset, and 101 in the test subset, each accompanied by their corresponding probability vectors and matrix.

**Training TSM in ME Detector.** We used annotated video segments with start and end timestamps to train TSM. The ResNet50 backbone network in TSM takes each frame of the query video as input and generates behavioral representations of equal length. These representations are integrated into several fixed-length (we set to 8) segments according to the given start and end timestamps, following the method in [4]. The FC network in TSM, a binary classifier, takes the integrated segments as input and predicts the ME probability. The training is conducted with the Momentum optimizer, using a momentum of 0.9 [5]. It starts with a learning rate of 0.005, which decays to 0.001 and 0.0001 after 20 epochs and 30 epochs, respectively. The batch size is set to 12, with data shuffled randomly. Training is conducted for 40 epochs with Cross-Entropy loss (CE), which computes the negative log-likelihood between the annotated and predicted segment labels as follows [6]:

$$\hat{Y}_{j}=\mathbf{FC}(\mathbf{concat}({\hat{\boldsymbol{b}}}_{1},{\hat{\boldsymbol{b}}}_{2},\ldots,{\hat{\boldsymbol{b}}}_{8};dim=0)),$$

$$L_{CE}=-\frac{1}{M}\sum_{j=1}^{M} \left[ Y_{j}\log\left( \hat{Y}_{j} \right)+\left( 1-Y_{j} \right)\log\left( 1-\hat{Y}_{j} \right) \right],$$

where **FC** is the single-layer fully connected classification network in TSM, **concat** is a function used to concatenate representation of several frames into a segment, ${\hat{\boldsymbol{b}}}_{i}$ is the behavioral representation of frame $i$, $\hat{Y}_{j}$ is the predicted label of segment $j$, $M$ is the number of segments, $Y_{j}$ indicates the annotation of segment $j$ (1 for ME, 0 for non-ME).

**Training BMN in ME Detector.** We trained TSM to generate behavioral representations for each frame. These representations, aligned with annotated probability vectors and matrix, were then used to train BMN. The training was conducted with the Adam optimizer, using default parameters as described in the original paper [7]. This starts with a learning rate of 0.001, which decays to 0.0001 and 0.00001 after 10 epochs and 15 epochs, respectively. The batch size was set to 12, with data shuffled randomly. Training was conducted for 20 epochs with the sum of two losses: a Weighted Binary Cross-Entropy loss (WBCE [8]) to compute differences between annotated and predicted start & end timestamp probability vectors, and a Proposal Evaluation Module loss (PEM [9]) to compute the regression and classification errors of segment probability matrices ($L_{reg}$, $L_{cls}$) as follows:

$$L_{WBCE}=-\frac{1}{M}\sum_{i=1}^{M} \left[ \boldsymbol{y}_{i}^{st}\log\left( {\hat{\boldsymbol{y}}}_{i}^{st} \right)+w\left( 1-\boldsymbol{y}_{i}^{st} \right)\log\left( 1-{\hat{\boldsymbol{y}}}_{i}^{st} \right)+\boldsymbol{y}_{i}^{ed}\log\left( {\hat{\boldsymbol{y}}}_{i}^{ed} \right)+w\left( 1-\boldsymbol{y}_{i}^{ed} \right)\log\left( 1-{\hat{\boldsymbol{y}}}_{i}^{ed} \right) \right],$$

Where $w$ is the weight assigned to negative samples to balance their contribution and is computed using the method [3], $M$ is the number of segments, $\boldsymbol{y}_{i}^{st} / {\hat{\boldsymbol{y}}}_{i}^{st}$ is the annotated/predicted probability vector for start timestamp, and $\boldsymbol{y}_{i}^{ed} / {\hat{\boldsymbol{y}}}_{i}^{ed}$ is the annotated/predicted probability for end timestamps.

$$L_{PEM}=10*L_{reg}+L_{cls},$$

Both $L_{reg}$ and $L_{cls}$ are used to quantify the difference between annotated and predicted segment probability of start-end timestamp pair forming an ME segment, following the method [3].

**Fine-tuning TSM in ME Refiner.** We used the predicted start and end timestamp probabilities, along with the segment probabilities (*i.e.*, the probability matrix), to compute the joint probability of each segment being an ME segment, following the method [3]. We then selected segments with joint probabilities greater than 0.01 as candidate ME segments. These segments were annotated as positive or negative samples (*i.e.*, 1 or 0) based on whether they overlap with the annotated ME segment, and are used for fine-tuning TSM in ME Refiner. The TSM is initialized by parameters from TSM in ME Detector and then trained by candidate ME segments using the same configurations as before.

**References**

[1] He KM, Zhang XY, Ren SQ, Sun J. Deep Residual Learning for Image Recognition. 2016 Ieee Conference on Computer Vision and Pattern Recognition (Cvpr) 2016: 770-778.

[2] Huang X, Cai Z. A review of video action recognition based on 3D convolution. Computers and Electrical Engineering 2023, 108.

[3] Tianwei Lin XL, Xin Li, Errui Ding, Shilei Wen. BMN Boundary-Matching Network for Temporal Action Proposal Generation. 2019.

[4] Lin J, Gan C, Wang K, Han S. TSM: Temporal Shift Module for Efficient and Scalable Video Understanding on Edge Devices. IEEE Trans Pattern Anal Mach Intell 2022, 44: 2760-2774.

[5] Sutskever I, Martens J, Dahl G, Hinton G. On the importance of initialization and momentum in deep learning. Proceedings of the 30th International Conference on International Conference on Machine Learning - Volume 28 2013: III–1139–III–1147.

[6] Domingos P. A Few Useful Things to Know About Machine Learning. Communications of the Acm 2012, 55: 78-87.

[7] Heo B, Chun S, Oh SJ, Han D, Yun S, Kim G*, et al.* AdamP: Slowing Down the Slowdown for Momentum Optimizers on Scale-invariant Weights. arXiv 2020, 2006.08217.

[8] Lin TY, Goyal P, Girshick R, He K, Dollár P. Focal Loss for Dense Object Detection. 2017 IEEE International Conference on Computer Vision (ICCV) 2017: 2999-3007.

[9] Lin T, Liu X, Li X, Ding E, Wen S. Bmn: Boundary-matching network for temporal action proposal generation. Proceedings of the IEEE/CVF international conference on computer vision 2019: 3889-3898.
